# Leveraging spectrum of graph sheaf Laplacian as a genome-architecture-aware measure of microbiome diversity

**DOI:** 10.64898/2026.03.10.710879

**Authors:** Nicolae Sapoval, Todd J. Treangen, Luay Nakhleh

**Affiliations:** Department of Computer Science, Rice University; Department of Bioengineering, Rice University; Ken Kennedy Institute, Rice University; Department of BioSciences, Rice University

**Keywords:** metagenomics, structural variants, horizontal gene transfer, microbiome diversity, genome graphs

## Abstract

**Motivation:** Measures of microbial diversity that can be derived directly from metagenomic sequencing data offer a valuable summary view of the underlying complex systems. Prior work has shown that both taxonomic composition and abundances that are captured by standard diversity measures (e.g., Shannon entropy), and structural variation within the metagenome due to gene duplications, losses and horizontal transfers (HGT), can correlate with the host’s health. However, there are no diversity measures available that simultaneously account for the genome architecture and taxonomic composition within the sample. Thus, in this work we propose the spectral energy of a graph sheaf Laplacian as such a measure, and justify its applicability through a simulation study and analysis of biological data.

**Results:** First, we describe a theoretical framework that allows us to combine the features of genome graphs with the taxonomic data. Then, we explore the sensitivity of the proposed diversity measure to genome rearrangements and HGT events in a simulation study. Finally, we explore applicability of our proposed measure to characterization of diversity of human gut metagenomes. We find our proposed measure to offer better discrimination between healthy controls and inflammatory bowel disease (IBD) patients’ samples (*n* = 403) in the cohorts analyzed.

**Availability and Implementation:** https://github.com/nsapoval/bd-gsl

## 1 Introduction

Metagenomics, the joint sequencing of uncultured microbial communities and the subsequent analysis of the sequencing data, allows in-depth investigations of microbial life and ecology and its impact on human health [Byrd et al., 2018, Xiao et al., 2020]. The two central questions of metagenomics are “Who is there?” and “What do they do?”. These are answered by taxonomic classification [Wood et al., 2019, Beghini et al., 2021, Blanco-Míguez et al., 2023] and functional characterization tools and analyses [Whittle et al., 2019, Yu et al., 2019, Moreno-Pino et al., 2020, Beghini et al., 2021]. However, since microbial communities are complex systems, these initial questions do not characterize the full spectrum of underlying complexity. Furthermore, previous work has shown that additional factors, such as genome architecture influenced by horizontal gene transfer (HGT) [Toft and Andersson, 2010, Treangen and Rocha, 2011] and structural variation (SV) [Durrant and Bhatt, 2019, Zhernakova et al., 2024], can be linked to health-related phenotypes in host-microbiome systems [Greenblum et al., 2015, Bonder et al., 2016, Zeevi et al., 2019, Zhernakova et al., 2024].

It is common to use complexity/diversity measures when studying properties of different ecological systems, and the microbiome is no exception. The classical measure of diversity for microbial data is the Shannon entropy [Shannon, 1948] of the relative abundance profile of the taxa that make up the biological sample. Previous work has shown that the entropy of the microbial sample can reflect host health or environmental conditions [Fan and Pedersen, 2021, Smith et al., 2019]. *However, the current paradigm based on taxonomic entropy for sample complexity is oblivious to the underlying genome architectures*. Given the prevalence of HGT in bacteria and its role as a driver of bacterial adaptation and response [Treangen and Rocha, 2011, Brito, 2021, Arnold et al., 2022, Bhaya et al., 2025], the insensitivity of diversity measures to this factor is problematic. Furthermore, SVs are emerging as key factors mediating host-microbiome interactions and host health [Zeevi et al., 2019, Zhernakova et al., 2024]. Previously, graph-based characterizations of sample complexity that track intra-sample SVs have been proposed [Ghurye et al., 2019, Balaji et al., 2022, Sapoval et al., 2024]. However, none of these methods take into account the taxonomy of the underlying genomic fragments.

In the current landscape of sample complexity/diversity characterization in metagenomics, a dichotomy exists between taxonomy-only and genome-graph-only methods (Fig. 1). However, there is ample evidence that both taxonomic composition [Fan and Pedersen, 2021, Smith et al., 2019] and genomic architecture [Durrant and Bhatt, 2019, Zeevi et al., 2019, Zhernakova et al., 2024] play key roles in emergent properties of the microbiome. In this work, we leverage a recent mathematical toolkit developed for modeling and characterizing data on graphs and formalized through the notion of a *sheaf on graph* [Friedman, 2015, *Hansen, 2020], in order to provide a first joint measure of metagenomic diversity that accounts for both taxonomic composition and genomic architecture of a sample (E*(*GSL*) in Fig. 1). We define the proposed diversity measure, formalized as the spectral energy of the sheaf [Davidson and Grinfeld, 2026] on a de Bruijn graph [De Bruijn, 1946]. We then provide results that show sensitivity of our proposed measure to genome rearrangements and HGT events in the underlying genomes. Finally, we demonstrate how our measure captures differences between samples derived from healthy controls and inflammatory bowel disease (IBD) patients across 403 metagenomic samples [Shaw et al., 2016, Franzosa et al., 2019, Smith et al., 2019]. We conclude the manuscript with a discussion of our results, some of the current limitations of our approach, and directions for future work.

**Figure 1.**
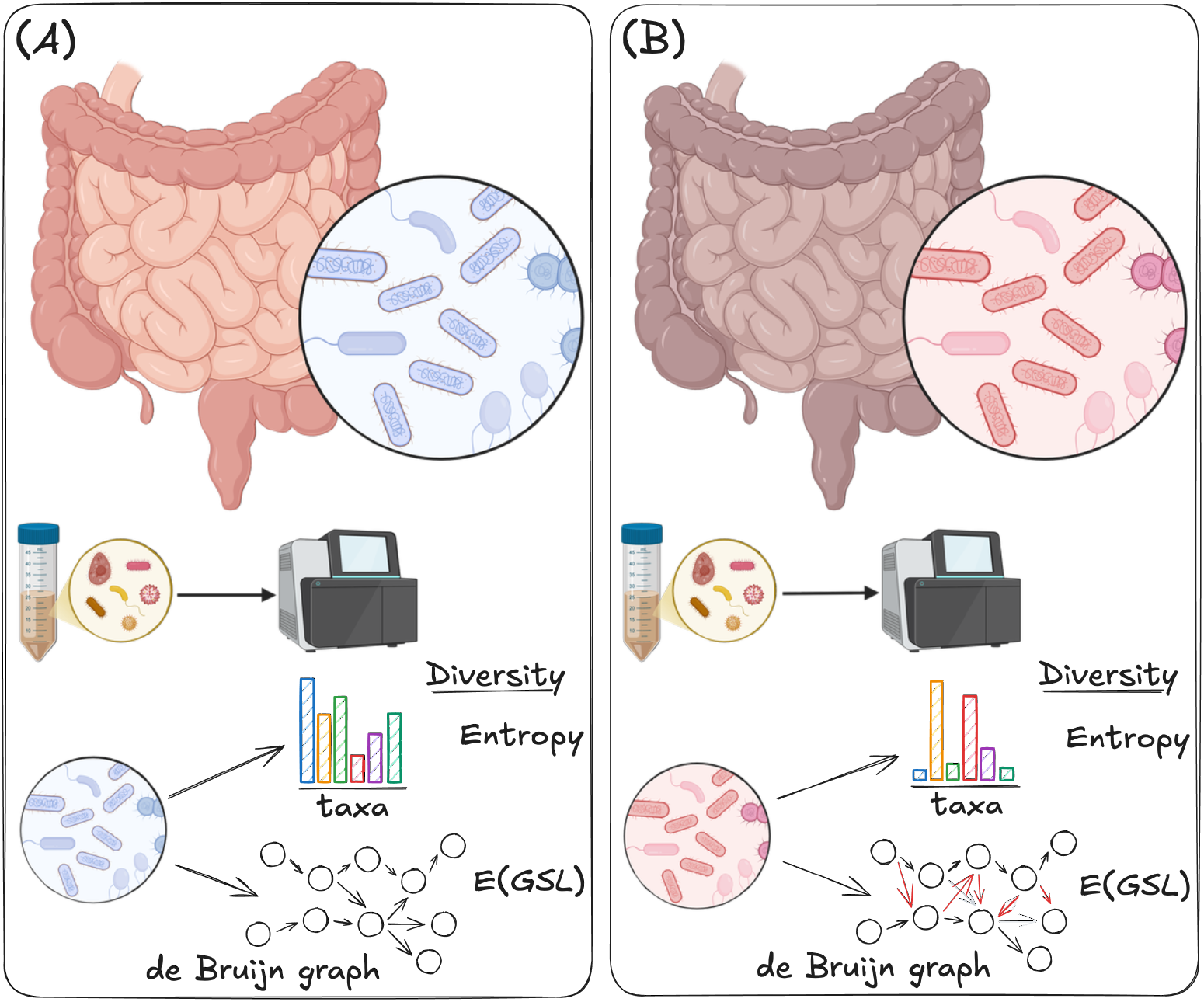
An overview of the metagenomic analysis of samples from a healthy donor (A) and a donor with an IBD condition (B). In both cases, a sample is preprocessed and then sequenced resulting in a collection of metagenomic sequencing reads that are used in the downstream analyses. Sample complexity/diversity is then quantified either by considering the taxonomic composition and relative taxon abundances or by analyzing the underlying de Bruijn graph structure. Our proposed approach merges these views through the spectral energy of the graph sheaf Laplacian.

## 2 Methods

### 2.1 Graph Sheaf Laplacian

#### Definition 1

(Graph sheaf). *Let G* = (*V, E*) *be a simple undirected graph. A sheaf ℱ on G (i*.*e*., *graph sheaf ℱ*_*G*_*) is a collection of vector spaces ℱ*(*v*) *and ℱ*(*e*) *for every vertex v ∈ V and edge e ∈ E respectively, together with a collection of* restriction maps *ℱ*_*v*⊴*e*_ : *ℱ*(*v*) *→ ℱ*(*e*) *defined for each incident pair v* ⊴ *e*.

For the rest of this manuscript we will assume that all vector spaces *ℱ* (*v*), *ℱ* (*e*) are copies of ℝ^*k*^ where the choice of *k* will depend on *v* and/or *e*. We will also assume that all restriction maps are unscaled projections.

#### Definition 2

(Cochains). *Let ℱ*_*G*_ *be a graph sheaf. We can then define the spaces of 0-dimensional and 1-dimensional* cochains *of ℱ*_*G*_ *as*

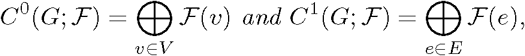

*respectively*.

Thus, a 0-cochain is an assignment of vectors to vertices of *G* and a 1-cochain is an assignment of vectors to the edges of *G*. Note that these assignments do not have to be compatible, i.e., they do not have to form a section.

#### Definition 3

(Coboundary map). *Let ℱ*_*G*_ *be a graph sheaf. Assume an arbitrary, fixed orientation of the edges of G and let* sign (*v*; *e*) ∈ {± 1} *denote the resulting incidence relationship. We define the* coboundary map *as*

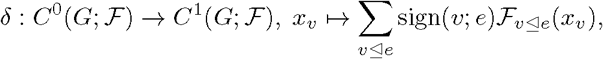

*extended by linearity to the entire C*^0^(*G*; *ℱ*).

We note that *δ* can be equivalently defined as (*δx*)_*e*_ = *ℱ*_*v*⊴*e*_(*x*_*v*_) − *ℱ*_*u*⊴*e*_ (*x*_*u*_) and thus constructed by iterating through all edges of *G*. Furthermore, for the purposes of the spectral analysis we are interested in, the fixed orientation of the graph does not matter.

#### Definition 4

(Graph sheaf Laplacian). *Let ℱ*_*G*_ *be a graph sheaf and let δ be its coboundary map. Then the symmetric positive semi-definite matrix L* = *δ*^*⊤*^*δ is the corresponding* graph sheaf Laplacian.

Finally, we define the energy of a graph sheaf Laplacian (GSL), which we use as the value of interest in the rest of the manuscript. We note that the spectrum of the Laplacian encodes strictly more information than its energy does, and hence can be used to further distinguish underlying structures within the data. However, we do not explore this direction in the present work.

#### Definition 5.

*Let ℱ*_*G*_ *be a graph sheaf and let L be its Laplacian. We define the energy of a Laplacian as* 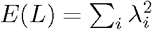, *where λ*_*i*_ *are the eigenvalues of L*.

Since, *E*(*L*) = Tr(*L*^2^) and equivalently it is the square of the Frobenius norm of *L*, we compute *E*(*L*) by computing the sum of squares of values in *L*. In order to avoid computation of the entire *L* at once, we do this in an iterative fashion using sparse representations of *δ*.

### 2.2 Formulation for sample diversity

For the rest of the manuscript we will fix the taxonomic rank of interest to be species. This choice doesn’t affect the mathematical formulas, but will be consistent for all empirical analyses performed. First, we will define the classical measure of microbial sample diversity given by the Shannon entropy.

#### Definition 6

(Shannon entropy). *Given a relative abundance profile* 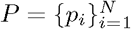 *for a sample containing N taxa. We define the Shannon entropy of the sample as:* 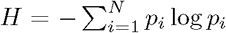. *We note that, by definition*, 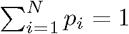

It is clear from the definition above that the entropy only depends on the relative abundances of taxa in the sample and does not account for the genomic architecture within it. In particular, if a microbial community undergoes active HGT without significant alteration in the abundances of constituent taxa, the entropy is insensitive to this change (Fig. 2).

**Figure 2.**
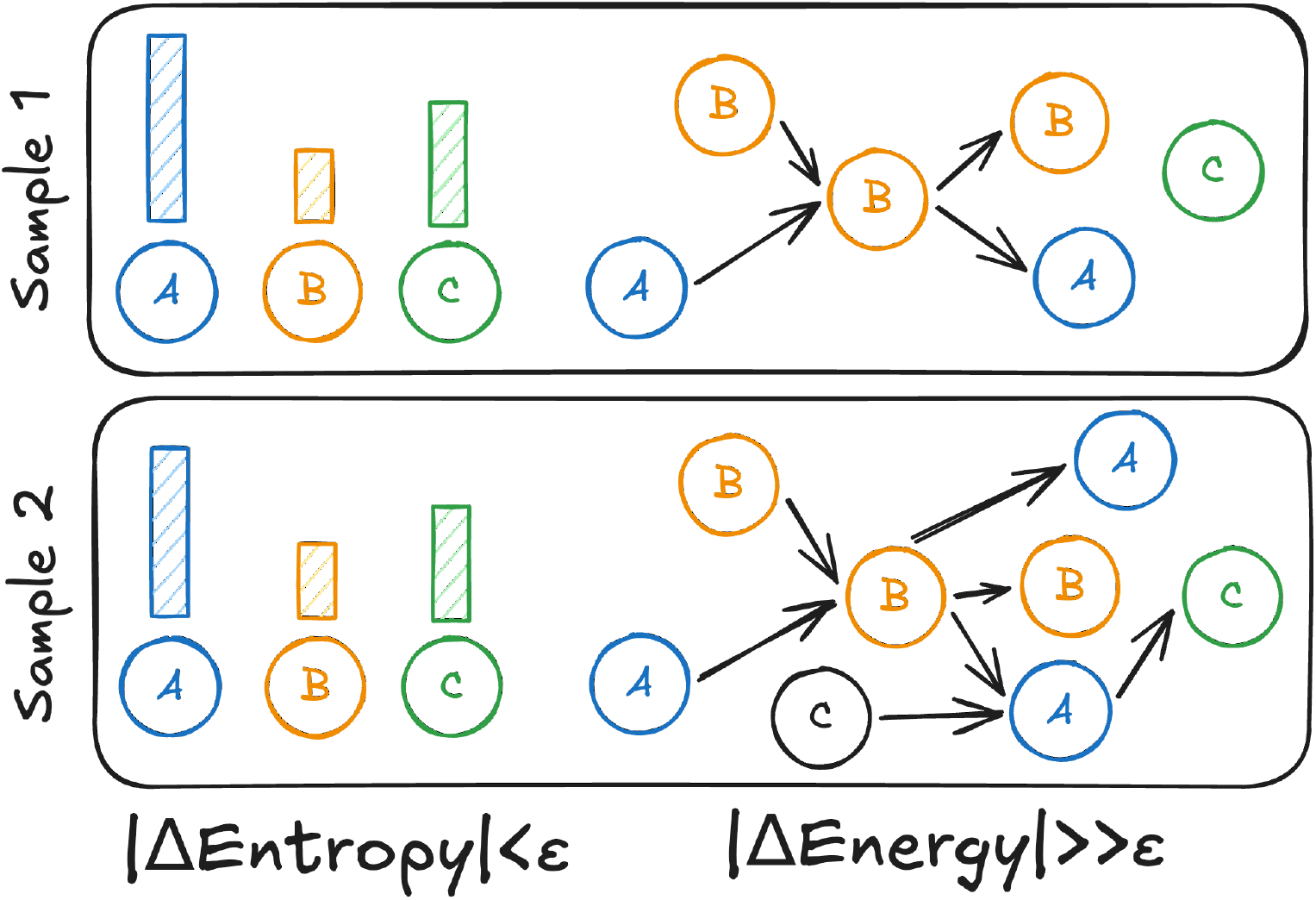
A schematic illustrating two samples with near identical relative abundance profiles (left), but distinct underlying genome architectures (right). Different species are indicated with colors and letters. The expected outcome is negligible difference in Shannon entropy of these two samples, yet a notable change in their respective GSL energies.

At the same time, a common first step towards metagenomic genome assembly is the construction of the de Bruijn graph [De Bruijn, 1946, Pevzner et al., 2001], which is equivalent to its compacted version in which non-branching *k*-mer paths are reduced to single vertices with corresponding strings that are called *unitigs* [Cracco and Tomescu, 2023]. Given that the unitigs themselves can be assigned taxonomic labels, while the underlying de Bruijn graph captures the information about (meta)genomic architecture of the sample, we propose the following definition for the GSL energy of a microbiome sample.

#### Definition 7

(GSL Energy). *Let G be the compacted de Bruijn graph of a metagenomic sample S. Let ℓ* : *V → {*0, 1*}*^*N*^ *be the taxonomic labeling for the vertices (unitigs) of G. We then define the graph sheaf, ℱ*_*G*_, *by letting ℱ*(*v*) = ℝ^*m*^, *where* 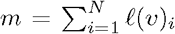. *Furthermore, let ℓ*_*∩*_(*u, v*) *denote the bit-wise AND of ℓ*(*v*) *and ℓ*(*u*). *We then let ℱ*(*e*) = ℝ^*t*^ *for e* = *{u, v}, where t* = *ℓ*_*∩*_(*u, v*). *Finally, the restriction maps ℱ* _*v*⊴*e*_ : ℝ^*m*^ → ℝ^*t*^ *are the natural projections. Let L be the GSL as defined in Def. 4, and let E be its energy as defined in Def. 5. We say that E is the GSL energy of the sample S*.

We emphasize that the definition above is introduced specifically to increase sensitivity of the analyses to within-sample changes in genome architecture (Fig. 2). However, unlike the Shannon entropy, GSL energy is not sensitive to changes in relative abundance that do not affect sample composition (i.e., the identity of constituent taxa) and genome architecture. Thus, the proposed measure should be thought of as a complementary way of characterizing sample diversity, rather than a replacement for the Shannon entropy.

### 2.3 Implementation details

#### Unitig construction

We use GGCAT [Cracco and Tomescu, 2023] v2.0.0 available through Bioconda for the unitig graph construction. We run GGCAT with a fixed *k*-mer size of 35 for all samples in this study. The value *k* = 35 was chosen based on the total number of unitigs, average length, and resulting graph connectivity after a parameter sweep for *k* ∈ {21, 25, 31, 35, 41, 45, 51} . We also note that *k* = 35 is a common choice for working with short-read paired-end data. We keep the default value of 2 for the minimum *k*-mer multiplicity, and require unitig link reporting (`-e` ) for the subsequent graph reconstruction.

#### Taxonomic classification

We use Kraken 2 [Wood et al., 2019] for unitig classification, with the default parameters and the `core nt` database (build date 10/15/2025). We do not filter any of the classifications. Lowest common ancestor (LCA) classifications above species level are treated as a simultaneous classification into all species under the LCA node that are present in the sample. For example, if some unitigs have been classified as species *A* and *B*, while another unitig has been classified as a genus *X* that contains *A, B*, and *C*, we will assign labels *A* and *B* to that unitig. Taxonomic classification for reads is performed with the same settings, except that no redistribution from LCA is done.

#### Graph sheaf construction

For each unitig in the compacted de Bruijn graph, we construct a bit vector of length equal to one plus the total number of species classifications. Then, the last bit of all vectors and the bits corresponding to taxa assigned to the unitig by Kraken 2 are set to 1. Thus, every node gets a space ℝ^*m*^ associated with it, where *m* is the number of non-zero bits in the corresponding vector. Subsequently, we compute the coboundary vector for each edge in the graph and construct the complete coboundary matrix *δ*. We compute and store *δ* as a compressed sparse row (CSR) matrix using `scipy.sparse`.

### 2.4 Details of the simulation

#### Data preparation

We selected 6,242 bacterial genomes with “complete” assembly level, annotations provided by NCBI RefSeq [Goldfarb et al., 2025], and marked as reference genomes. We then downloaded genomic FASTA and genome annotations using NCBI datasets CLI [O’Leary et al., 2024]. Additionally, to reduce the impact of closely related taxa, mismatches between species and strain taxonomic IDs, and candidate species on the taxonomic classification performed, we excluded from the simulation all *E. coli* and *Shigella* genomes, as well as any *Candidatus sp*.

#### Simulated genome rearrangements

In order to investigate sensitivity of the GSL energy to perturbations within a genome of a single bacterial species, we performed a genome rearrangement experiment. Specifically, we sampled a genome from the downloaded data described above and produced shuffled copies of the genome with gene order changed, and some of the genes replaced by their reverse complements, simulating inversions. Then short paired-end reads were simulated from three mixtures (original genome, original genome +1 genome with rearrangements, and original genome +10 genomes with rearrangements) of genomes using ART [Huang et al., 2012]. We selected five *Synechococcus* species and five other species as the starting genomes for this experiment. The choice of *Synechococcus* was motivated by empirical evidence of extensive genome rearrangements in this genus [Birzu et al., 2025, Bhaya et al., 2025]. In all experiments, 100,000 reads-per-sample were simulated with a uniform abundance profile.

#### Simulated HGT metagenomes

Next, we simulated non-HGT and HGT bacterial communities by randomly sampling genomes from our downloaded dataset and simulating HGT events with HGTSim [Song et al., 2017]. We considered two dataset sizes: minimal – consisting of 2-3 genomes with a single HGT donor, and small – consisting of 10 genomes with 2-3 donors and 8 or 7 recipients, respectively. For the minimal dataset, we had on average 4 HGT events per sample, and for the small dataset we had an average of 14.5 HGT events per sample. Additionally, for both datasets, we considered medium (100-170 HGT events per small sample) and high (550-860 HGT events per small sample) HGT regimes. We explored both perfectly conserved HGT events, where the recipient received the exact copy of the donor gene, and HGT with mutation using the mixed mutation model from HGTSim. For the minimal dataset, we simulated 100,000 short paired-end reads for each sample, and for small dataset we simulated 500,000 reads for each sample. All reads were simulated with ART, and we considered both uniform and log-normal relative abundance profiles. Finally, in order to explore the impact of HGT alone, we also considered a set of experiments that use several identical genome copies, instead of simulated reads for the initial unitig construction.

## 3 Results

Results are organized into two main subsections. In the first subsection, we investigate the sensitivity of our proposed measure to changes in the genomic architecture of the sample constituents on simulated data. In the second subsection, we analyze a total of 403 short-read human gut metagenomes coming from healthy controls (HC) and patients with ulcerative colitis (UC) [Shaw et al., 2016, Smith et al., 2019] and Crohn’s disease (CD) [Franzosa et al., 2019, Smith et al., 2019]. We explore differences in entropy for these metagenomes, computed from both prior analyses using MetPhlAn and re-computed with Kraken 2, and our GSL energy calculations.

### 3.1 Simulated data

#### 3.1.1 Sensitivity to genome rearrangements

First, we note that in the genome rearrangement simulations the total GSL energy varies between the base genome samples and the samples that contain a mixture of rearranged genomes (Fig. 3). As expected, the entropy does not vary significantly, since all reads come from the genome of the same bacterial taxon and the genomes differ only in their organization.

**Figure 3.**
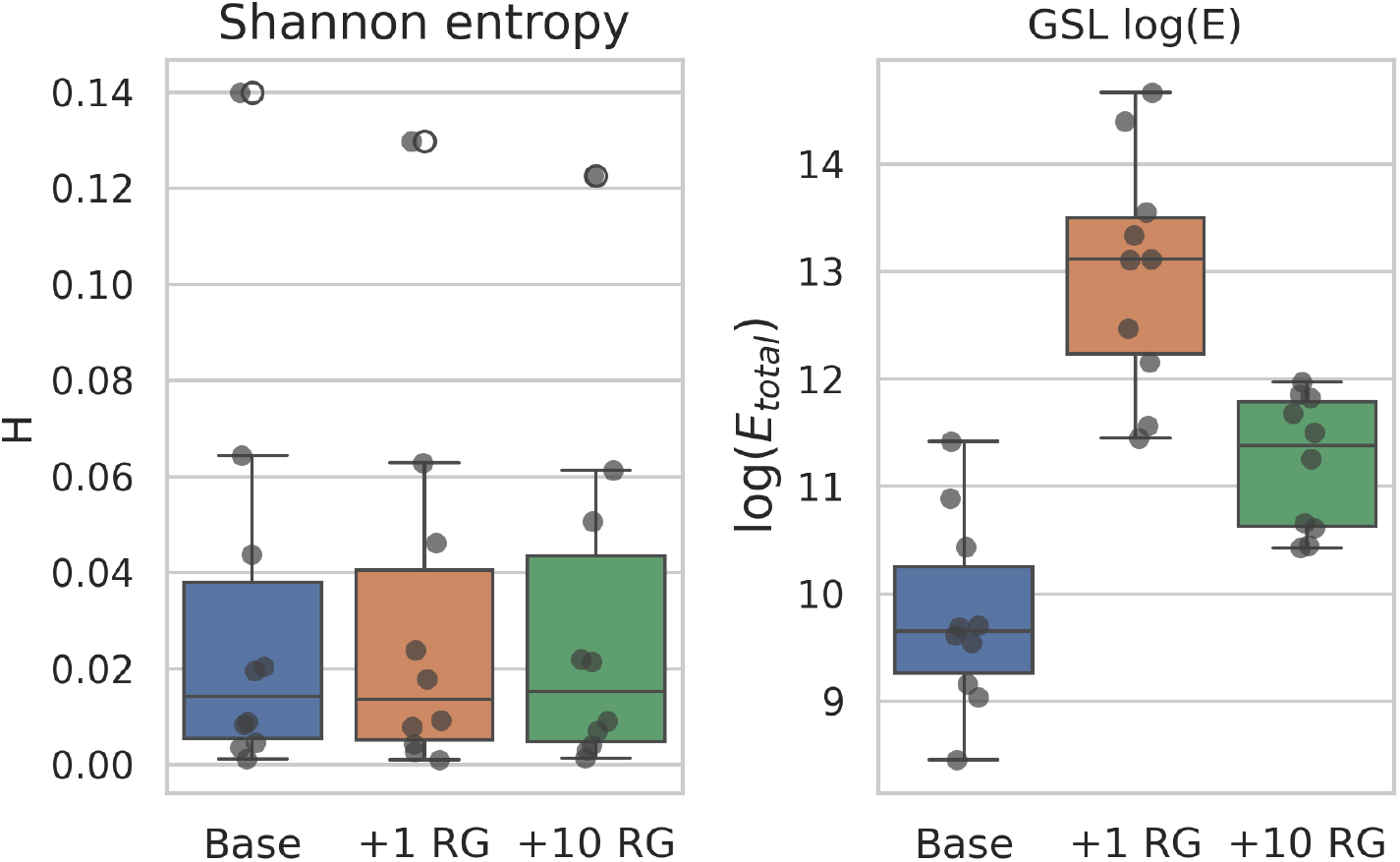
Shannon entropy (left panel) and the log of the GSL energy for the genome rearrangement experiments. Blue box shows distribution of values for the sample only containing the base genome, orange is a mixture of the base and 1 rearranged genome, and green is a mixture of the base and 10 rearranged version of the genome.

Notably, adding a single genome with rearrangements achieves the highest energy values, while adding 10 different genomes with rearrangements achieves higher energy than the base case, but lower than that of a simple mixture (Fig. 3). This is a direct reflection of the change in the underlying graph, as with more genomes the number of connected components increases, but the size of the largest component decreases (Supp. Fig. 10). Also, in the case of the genome rearrangement simulation, we note that the structural variation increases GSL energy (Supp. Fig. 11).

#### 3.1.2 Sensitivity to HGT events

Next, we evaluated whether the GSL energy of the sample would vary in the presence of HGT. First, we considered the case where unitigs are constructed directly from the genomes. We note that the overall magnitude of the differences in the total energy between the non-HGT and HGT samples is small (Fig. 4 and 5; left). However, similarly to the case of genome rearrangements, the direction of the change is largely consistent, with all samples under the minimal scenario showing minor positive deltas (Fig. 4), and all but 2 samples in the small scenario showing moderate positive deltas (Fig. 5).

**Figure 4.**
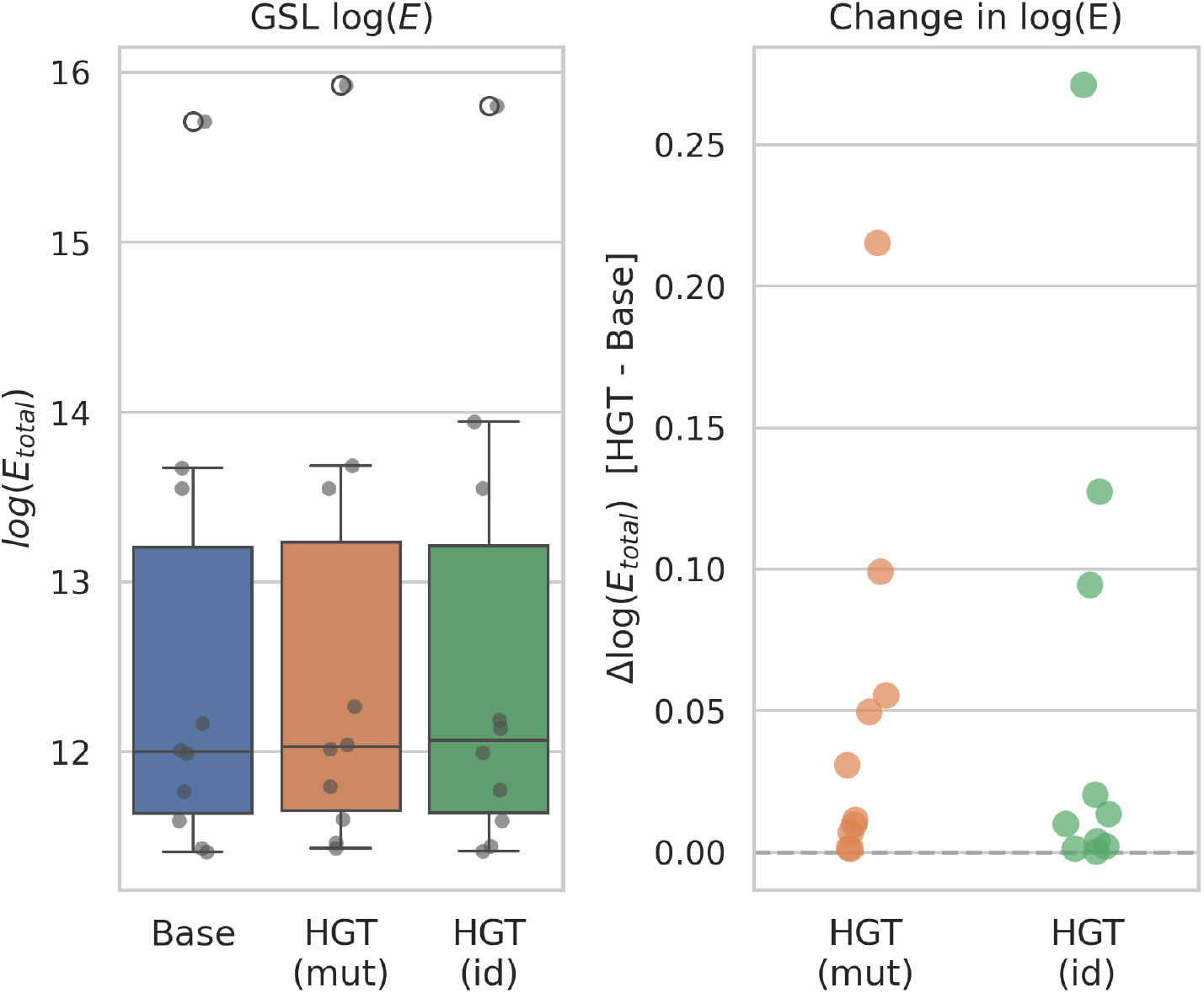
Logarithm of the GSL energy (left) and the change in the log of the GSL energy from baseline sample (right) for the minimal dataset. Orange shows the HGT simulation that mutates transferred gene, while the green shows results for the transfer of an identical gene copy.

**Figure 5.**
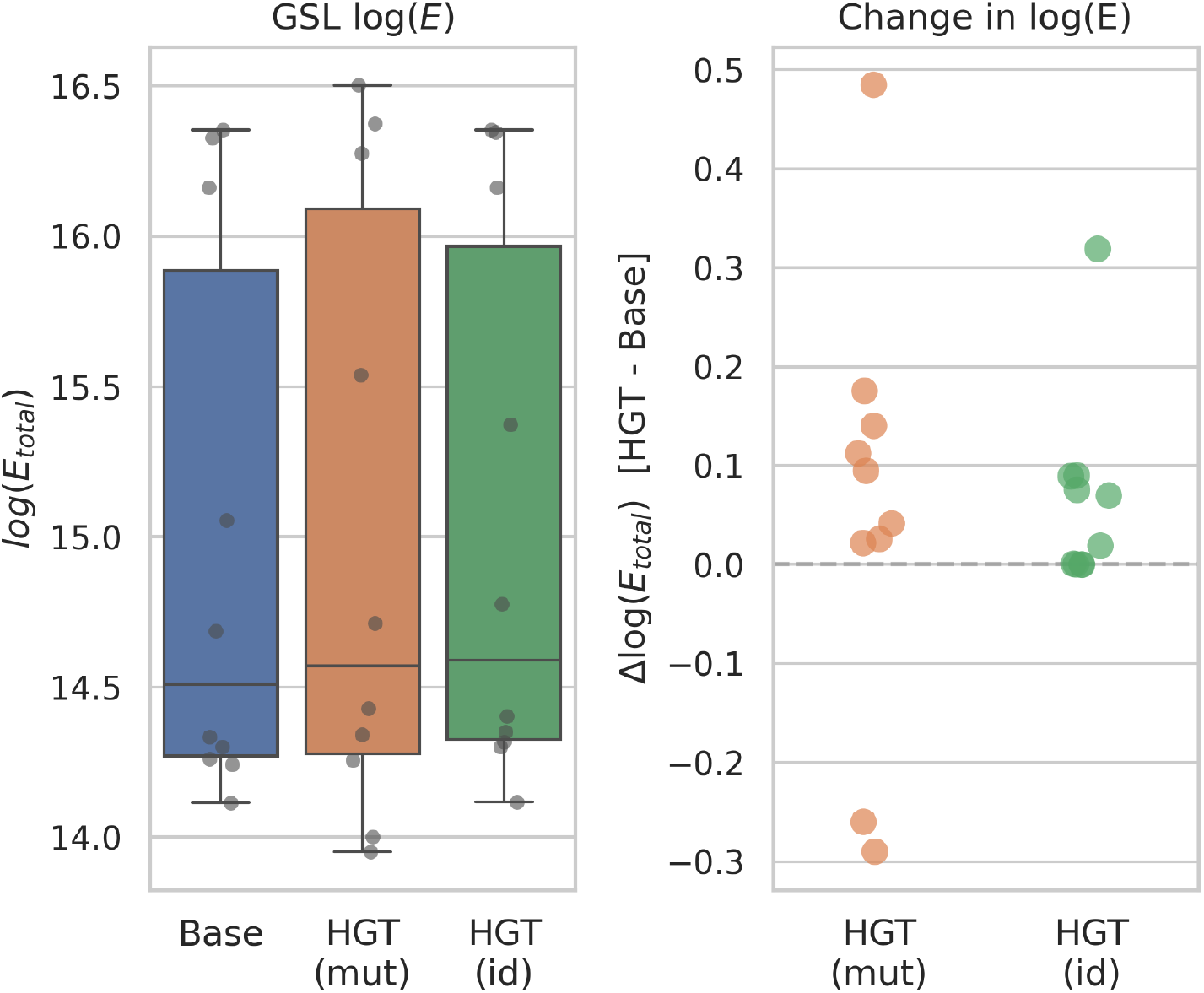
Logarithm of the GSL energy (left) and the change in the log of the GSL energy from baseline sample (right) for the small dataset. Orange shows the HGT simulation that mutates transferred gene, while the green shows results for the transfer of an identical gene copy.

We also evaluated the impact of HGT on the GSL energy on simulated metagenomic reads. We noted that the overall differences in GSL energy are more pronounced than in the case of the unitigs constructed from genomes (Fig. 6, 7). Furthermore, as expected, there is nearly no difference in the per sample entropy between baseline samples with no HGT and the ones that contain HGT events. This observation is consistent across abundance conditions and the type of transferred genes (mutated or identical).

**Figure 6.**
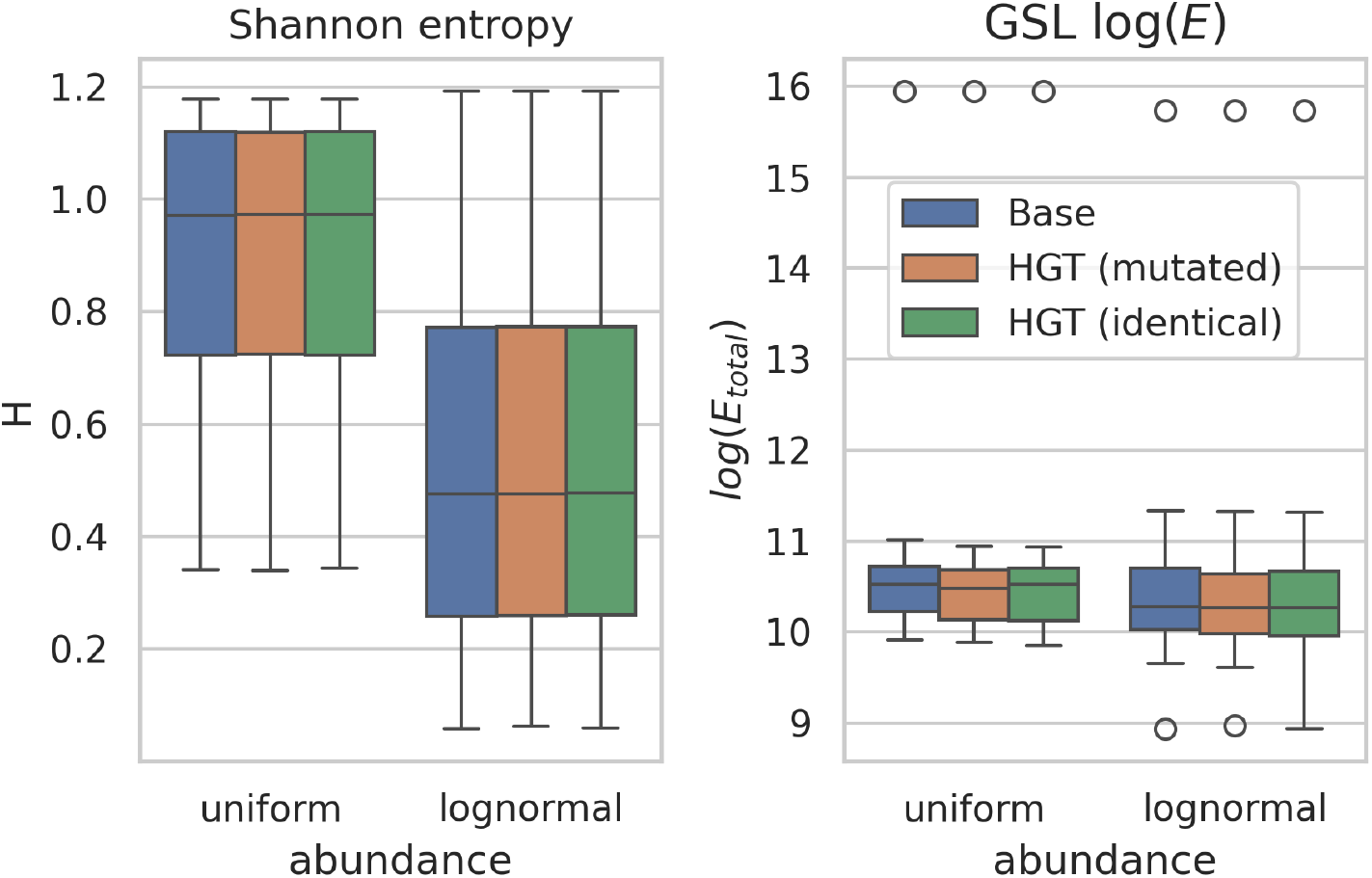
Shannon entropy (left) and the log of GSL energy (right) for the minimal samples with uniform and lognormal abundances. Colors indicate the baseline sample (blue), HGT with mutated transferred sequences (orange), and HGT with identical transfers (green).

**Figure 7.**
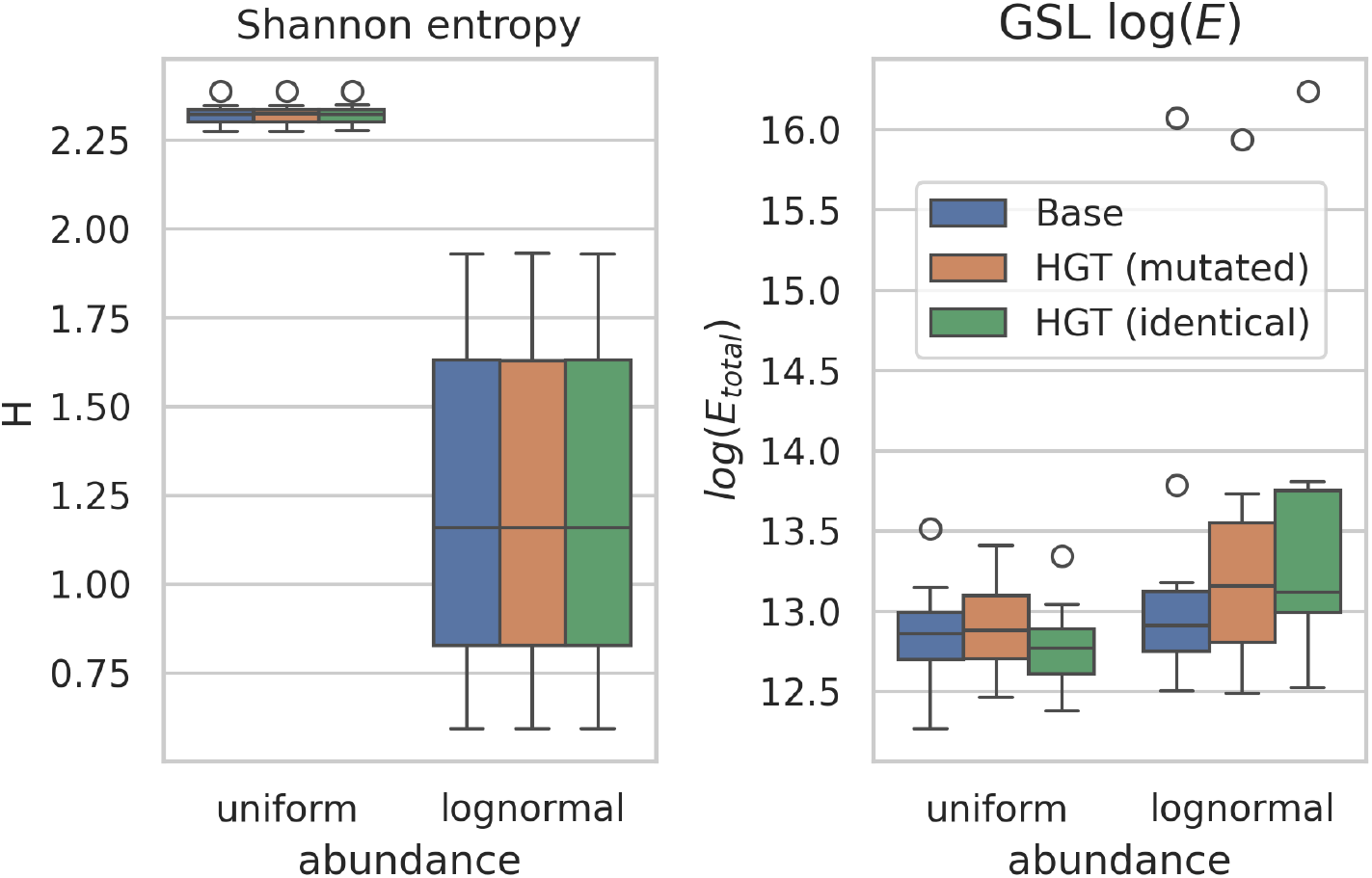
Shannon entropy (left) and the log of GSL energy (right) for the small samples with uniform and lognormal abundances. Colors indicate the baseline sample (blue), HGT with mutated transferred sequences (orange), and HGT with identical transfers (green).

Finally, we investigated whether the number of HGT events per sample affects the GSL energy. We noted that Shannon entropy remains insensitive to the HGT events regardless of their number, as expected (Supp. Fig. 12, 13). The GSL energy does increase overall for the higher rate of HGT events, and the gap between the baseline and HGT sample slightly widens with the number of events increasing (Supp. Fig. 12, 13).

### 3.2 Human gut metagenomes

First, we explored the Shannon entropies and GSL energies of the human gut metagenomic samples from the three studies (Fig. 8). We note that on the biological samples, Shannon entropy from both the prior reported analyses with MetaPhlAn and independent re-evaluation with Kraken 2 has discriminative power to separate healthy control samples from the ones with IBD conditions (Fig. 8). Similarly the GSL energy is also capable of separating these conditions, and leads to a tighter clustering and hence a better separation between the conditions (Fig. 8).

**Figure 8.**
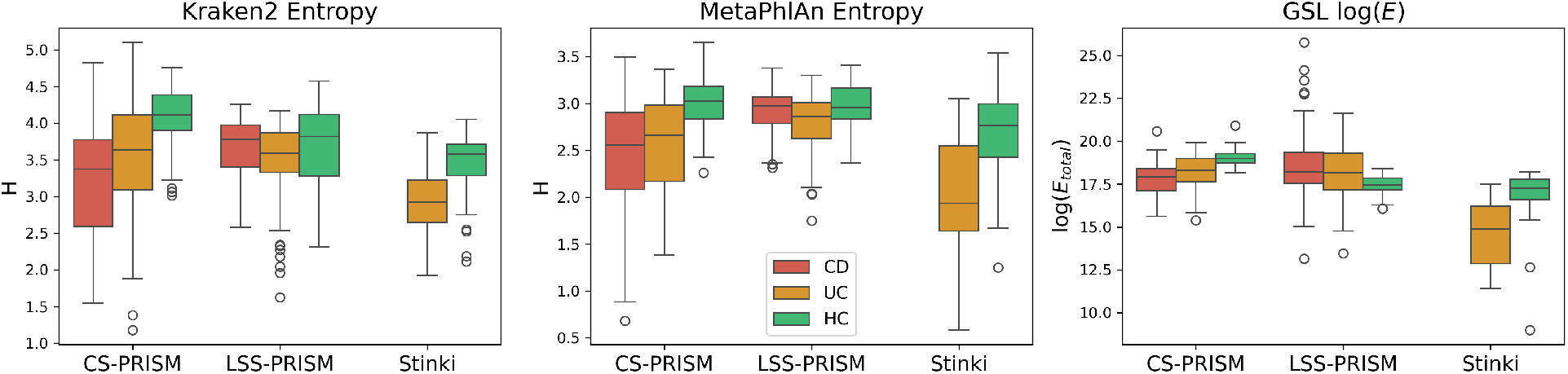
Distributions of the Shannon entropy and the log of GSL energy for the samples in CS-PRISM (*n* = 150), LSS-PRISM (*n* = 153), and Stinki (*n* = 87) cohorts. Colors indicate the patient status as a healthy control (HC, green), Crohn’s disease (CD, red) patient, or ulcerative colitis (UC, orange) patient.

In particular, for the CS-PRISM study (*n* = 151) all diversity metrics achieve a *p*-value *<* 0.05 for the Mann-Whitney U test for HC vs UC (K2: 0.0011, MPA: 0.0006, GSL: 0.0001) and HC vs CD (all *<* 10^−4^) populations (Supp. Tab. 1). For the LSS-PRISM study (*n* = 165) only GSL energy achieves *p*-values below *<* 0.05: 0.0401 for the HC vs UC populations and 0.0004 for the HC vs CD population (Supp. Tab. 1). For Stinki study (*n* = 87) all metrics achieve *p*-values *<* 10^−4^ (Supp. Tab. 1). These observations are matched by the AUC ROC values for classification between HC and UC, and HC and CD (Fig. 9). For all studies, GSL energy consistently offers best separation between the conditions (Fig. 9).

**Figure 9.**
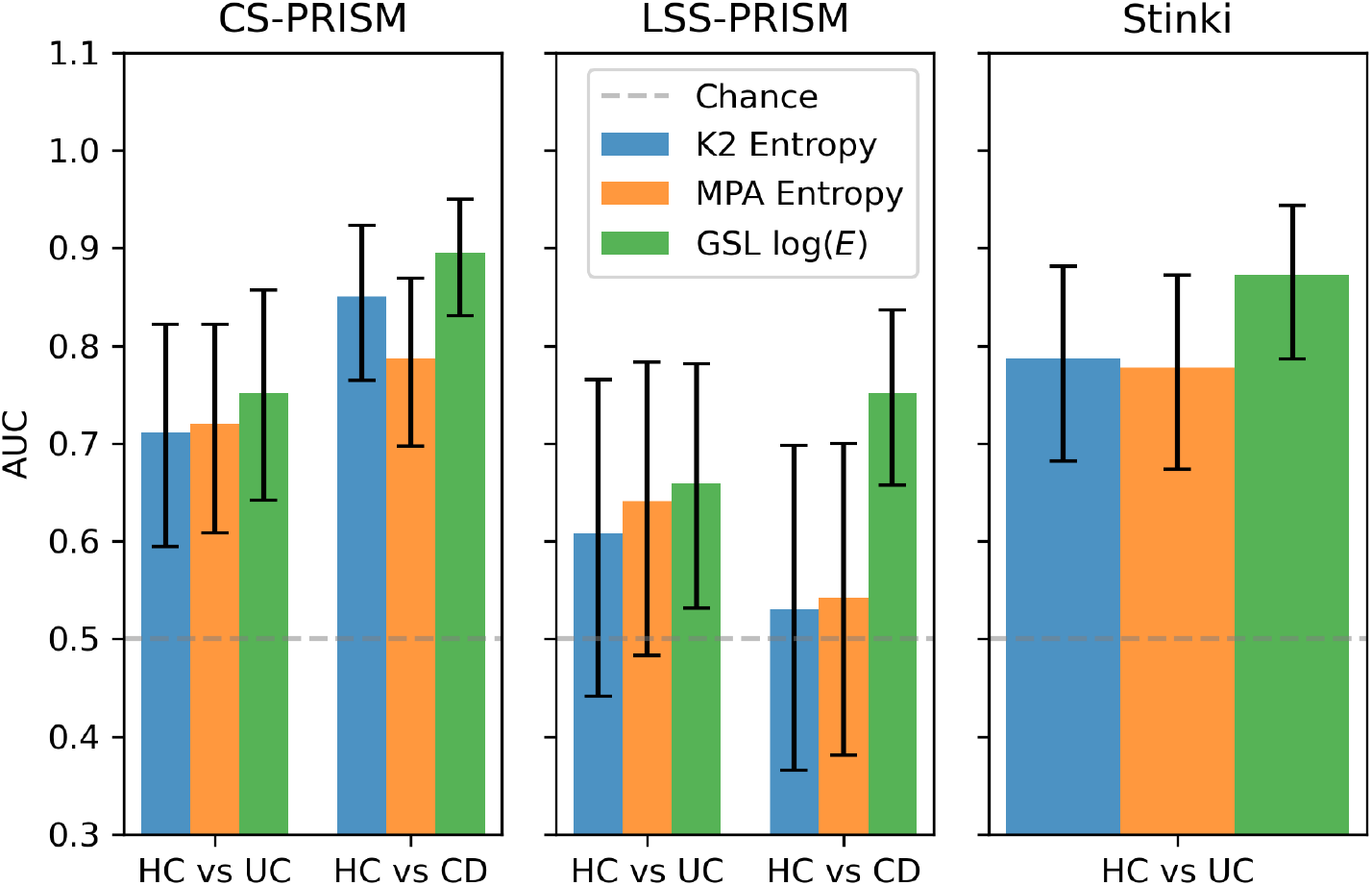
Area under the receiver operator characteristic curve (AUC ROC) for classification into HC and UC or CD status. Dashed gray line indicates a random choice baseline, and color bars indicate metrics (blue: K2 entropy, orange: MPA entropy, and green: GSL energy). Error bars indicate the 95% CI obtained from subsampling bootstrap.

The individual ROC curves (Supp. Fig. 14, 16) show that GSL energy achieves highest TPR for the low FPR thresholds. Additionally, we investigated the correlation between GSL energy and entropy computed from the biological samples, and found weak and no correlation for HC, and weak to moderate correlation for UC and CD samples (Supp. Fig. 15, 17).

**Figure 10.**
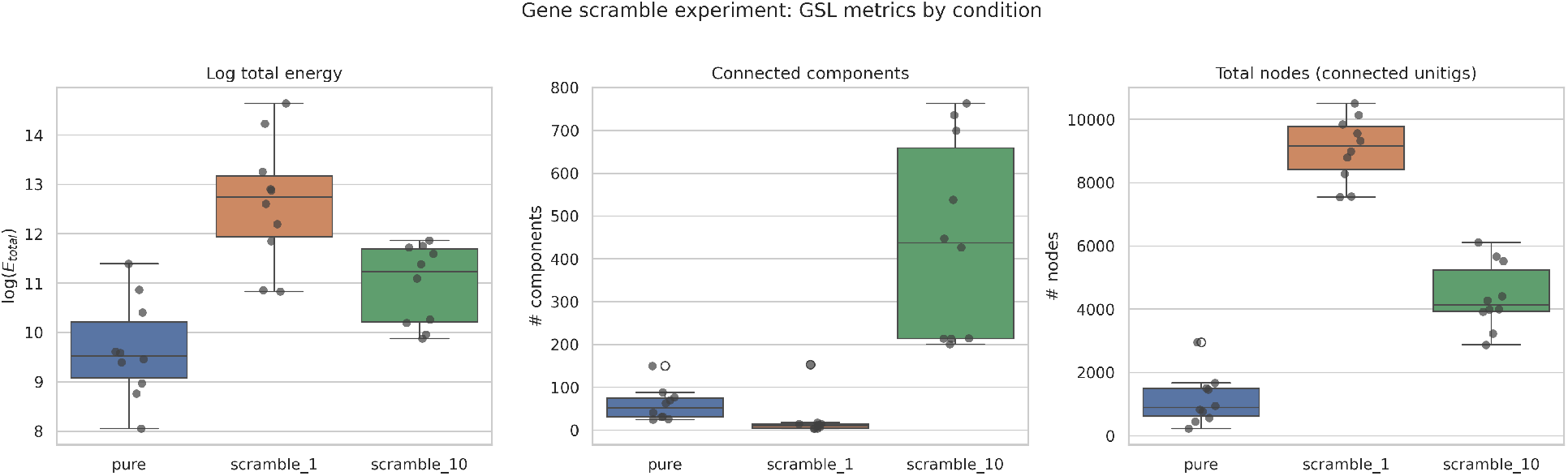
Logarithm of the total energy (left), number of connected components (middle), and total number of nodes (right) for a the single base genome (blue), a mixture of base genome and its rearranged version (orange), and mixture of base genome and 10 different rearranged copies (green).

**Figure 11.**
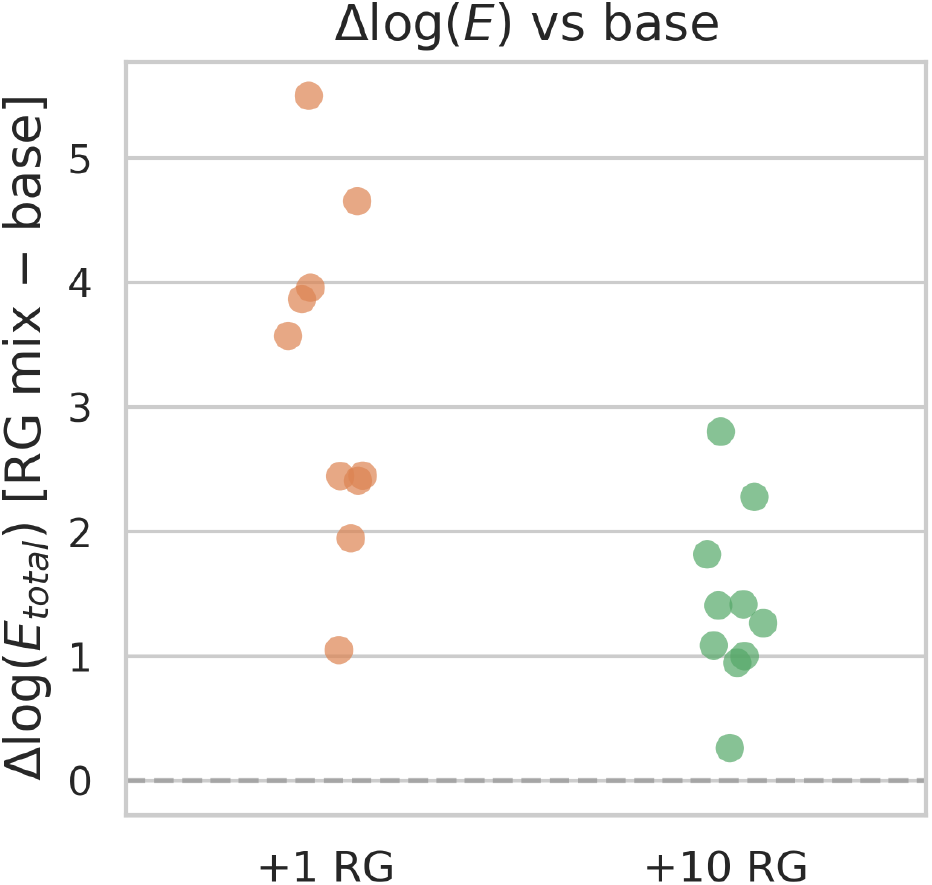
Change in the GSL energy between the base samples containing reads from a single genome to mixtures with 1 and 10 genomes with rearrangements. All observed changes are strictly positive.

**Figure 12.**
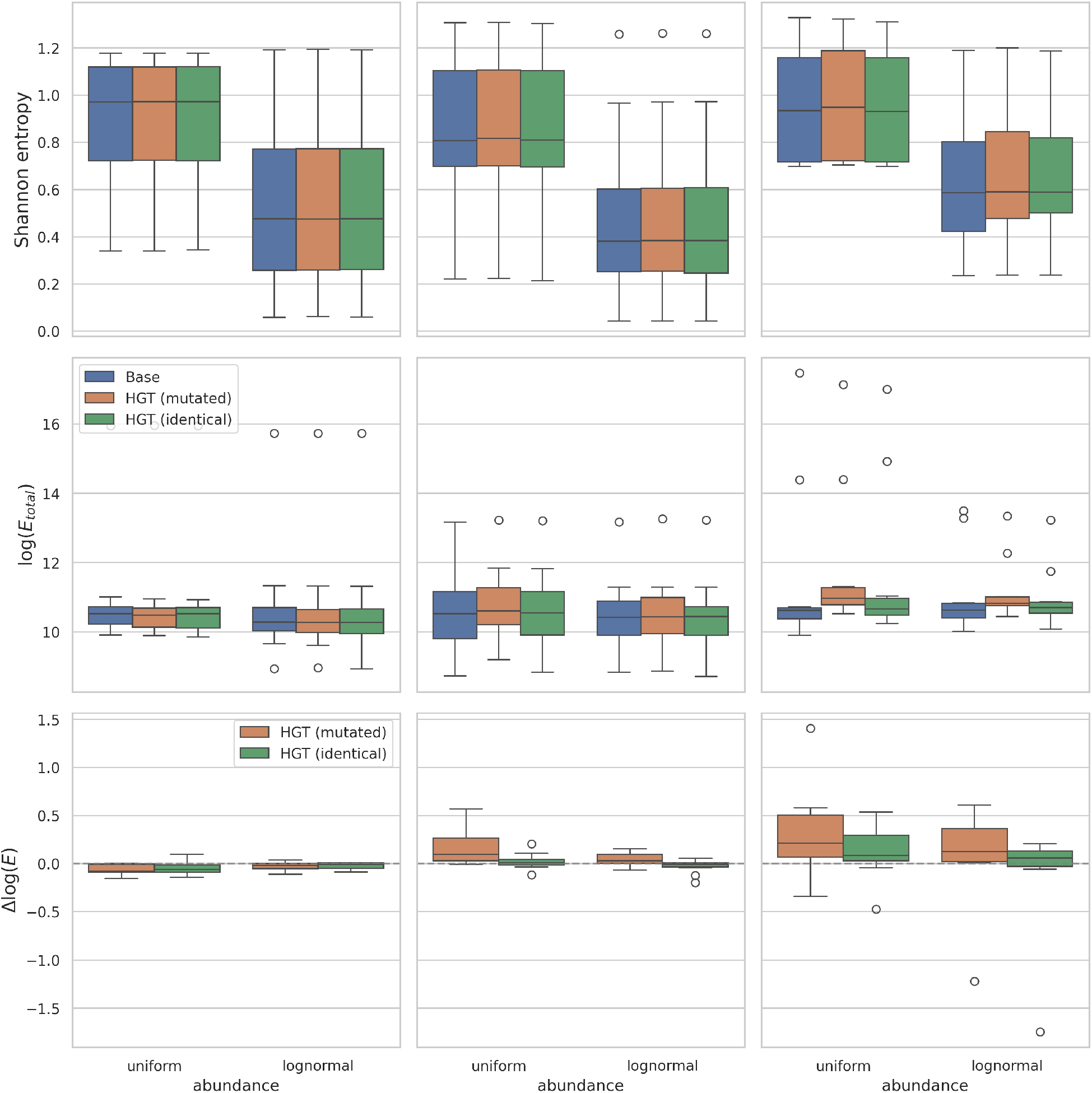
Shannon entropy (top row), logarithm of the GSL energy (middle row), and change in the logarithm of the GSL energy between the base and HGT samples (bottom row). Each column corresponds to the amount of genes affected by HGT: left is the baseline, middle is 10x the rate of the baseline, and right is 50x the rate of the baseline. All samples are on the minimal dataset consisting of 2-3 genomes.

**Figure 13.**
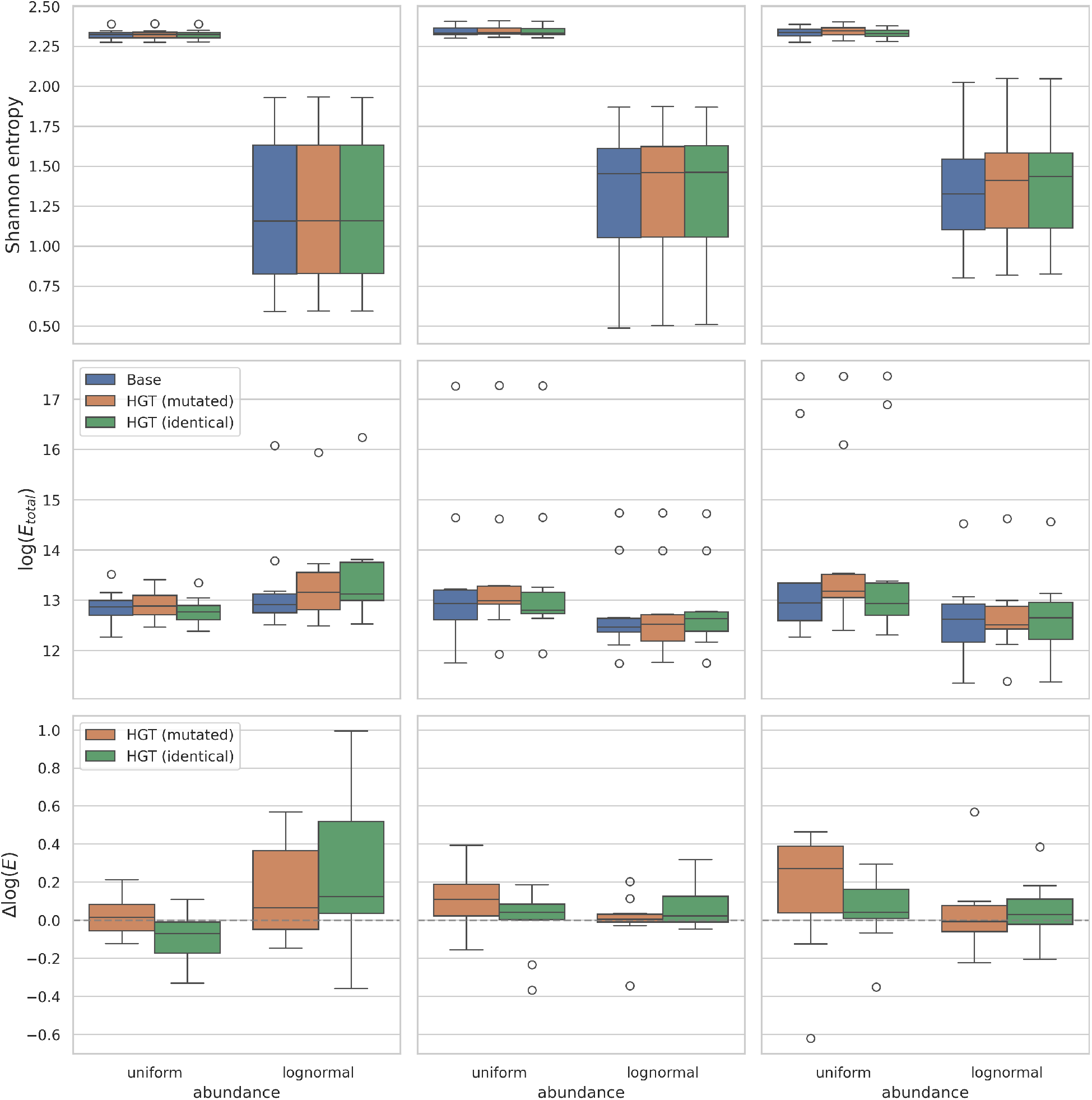
Shannon entropy (top row), logarithm of the GSL energy (middle row), and change in the logarithm of the GSL energy between the base and HGT samples (bottom row). Each column corresponds to the amount of genes affected by HGT: left is the baseline, middle is 10x the rate of the baseline, and right is 50x the rate of the baseline. All samples are on the small dataset consisting of 10 genomes.

**Figure 14.**
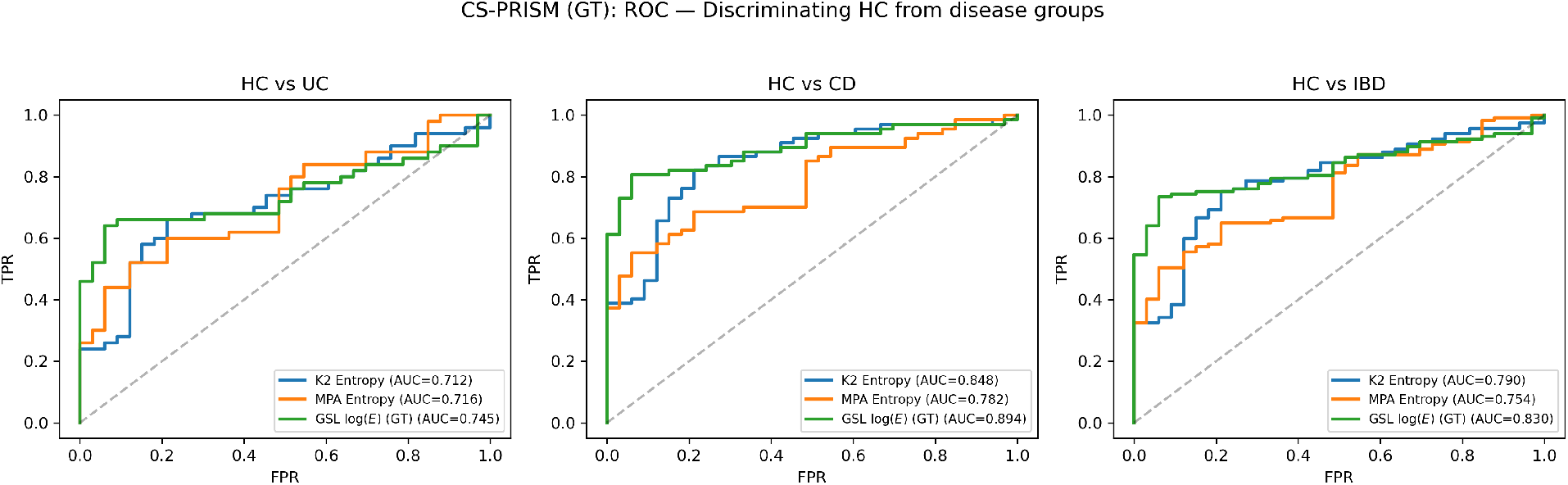
Receiver operator characteristic curves for separating power between UC, CD, and combined (IBD) conditions and healthy controls (HC). Evaluation done for the samples in CS-PRISM study cohort.

**Figure 15.**
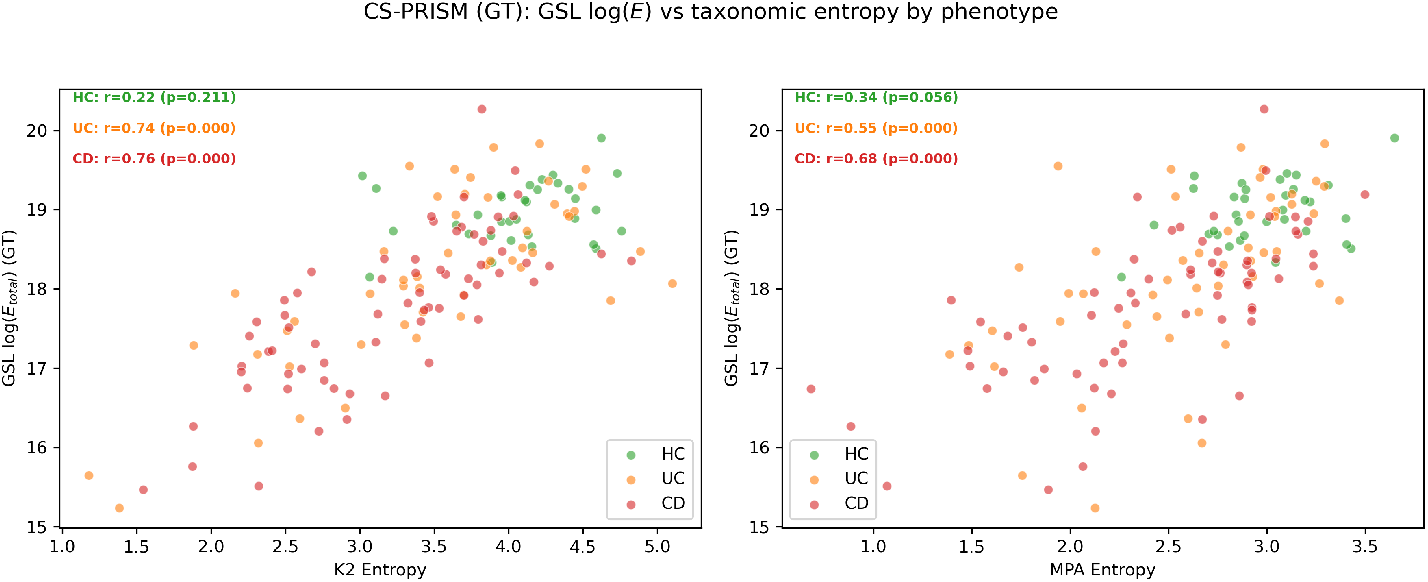
Per group (HC, UC, CD) correlation between Shannon entropy (from K2 and MPA classifications) and GSL energy for the CS-PRISM cohort samples.

**Figure 16.**
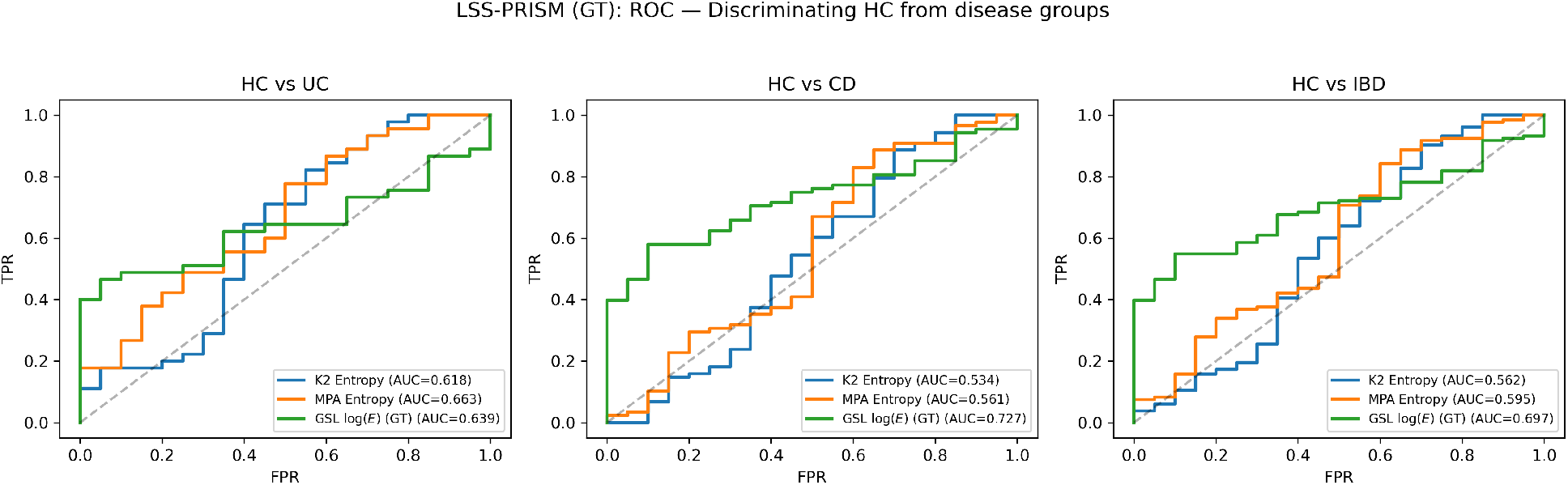
Receiver operator characteristic curves for separating power between UC, CD, and combined (IBD) conditions and healthy controls (HC). Evaluation done for the samples in LSS-PRISM study cohort.

**Figure 17.**
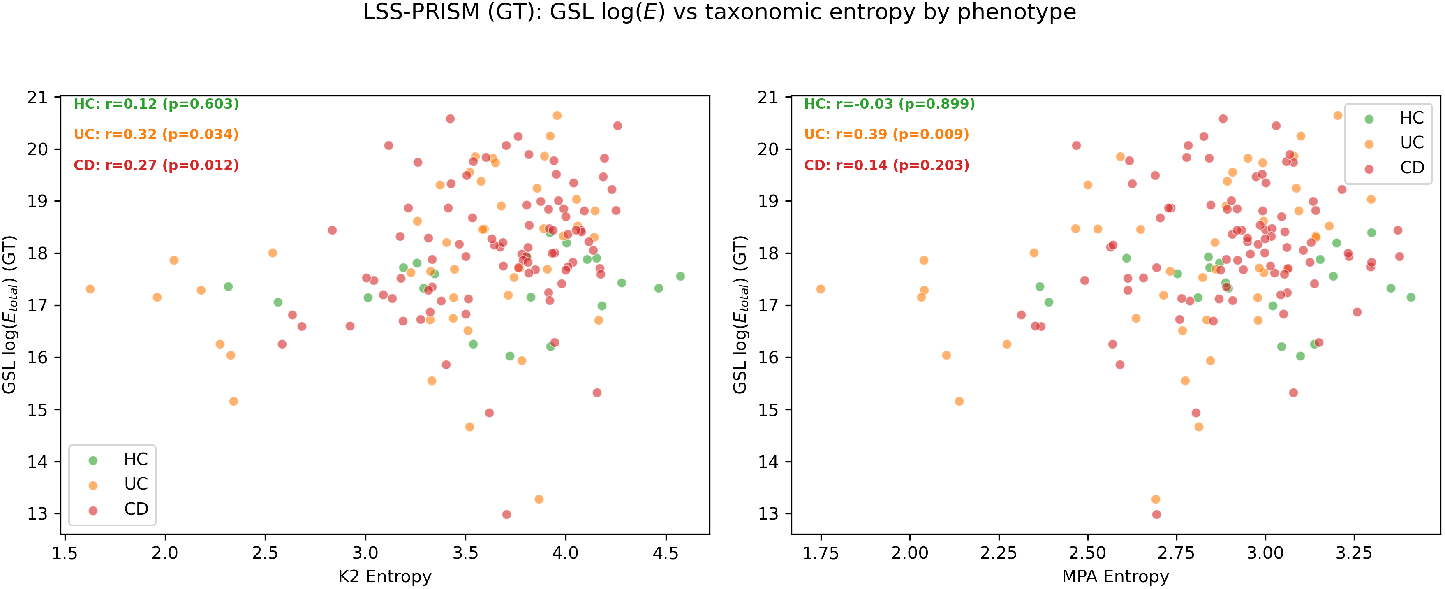
Per group (HC, UC, CD) correlation between Shannon entropy (from K2 and MPA classifications) and GSL energy for the LSS-PRISM cohort samples.

### 3.3 Computational performance

We measured resource usage for the CS-PRISM and LSS-PRISM samples (*n* = 303). All computations were performed on the 96-core Intel Xeon Platinum 8468 CPUs available through the NOTS cluster. All runs had access to up to 192 GB of RAM. CS-PRISM runs took a total of 136h 43m of wall clock time (CPU: 201h 27m) to complete, with the per sample average wall time of 54m 19s (CPU: 1h 20m). Fastest sample completed in 4m 28s, and the longest took 4h 40m (CPU: 7h 56m). LSS-PRISM runs took a total of 58h 51m of wall clock time (CPU: 60h 11m) to complete, with the per sample average wall time of 21m 24s (CPU: 21m 53s). Fastest sample completed in 32s, and the longest took 1h 17m (CPU: 2h 23m). Peak RAM usage was 13.06 GB achieved by one of the CS-PRISM samples, and the average RAM usage was 0.95 GB for CS-PRISM samples, and 0.17 GB for the LSS-PRISM samples.

## 4 Discussion

Our results show that using the energy of the GSL derived from taxonomic classifications over a metagenomic de Bruijn graph leads to a measure of microbiome diversity that is sensitive to both taxonomic composition and genomic architecture. Furthermore, the exploratory results on real human gut metagenomes indicate a strong discriminative potential for the GSL energy, supporting it as a candidate for linking microbiota genotype dynamics to host phenotype outcomes. Additionally, lack of strong correlation with the Shannon entropy, indicates that GSL energy can be used as a complementary characteristic of microbiome diversity. Together these observations indicate that our proposed diversity measure that accounts for both genomic and taxonomic complexity has greater resolution than a purely taxonomic alternative, and is meaningfully correlated with the phenotypic traits of interest.

We also recognize that our proposed measure has several limitations. First, as mentioned in Sec. 2.2, it is unable to distinguish between the samples with identical genomic composition, but varied genome abundance. However, since sheaves on graphs present a general framework for modeling of data on graphs, we believe that it is possible to extend our construction to account for abundance (e.g., by using read-to-unitig relationships and lifting read classifications). The key challenge we envision for this direction comes from the desired property of having a totally disconnected graph agreeing with the Shannon entropy calculation. We plan to explore the required mathematical constructions and the resulting complexity metrics in our future work. The second limitation is the lack of immediate interpretability of the change in the GSL energy. In particular, the directionality of the change between two samples is dependent on multiple factors. Hence, while two samples from different conditions will have distinct energies, we cannot always classify the sample based only on this value. Third, the current implementation of the GSL analysis is not heavily optimized and hence has limited support for very large graphs. Given that the Laplacian is a structured matrix, additional improvements to runtime and memory scalability might be achievable with optimized subroutines.

In conclusion, we introduced a novel measure of microbiome sample diversity that simultaneously accounts for genome architecture (SVs, HGT events) and taxonomic composition of the sample. We demonstrated its utility in simulated data settings under varying scenarios. Finally, we showed that this measure is sensitive and achieves strong separation between controls and condition samples of human gut metagenomes.

## 5 Acknowledgments

This work was in part supported by the NSF grants DMS/NIGMS-2153704 and DBI-2030604, and the NOTS cluster operated by Rice University’s Center for Research Computing (CRC). TJT was supported in part by NIH NIAID P01-AI152999, and NSF awards IIS-2239114, EF-2126387.

## A Supplemental figures and tables

**Table 1:** Mann–Whitney U and Brunner–Munzel tests for healthy controls vs. IBD group comparisons across the cohorts.

| Study | Comparison | Metric | $n_{\text{HC}}$ | $n_{\text{dis}}$ | $U$ | $p_{\text{MW}}$ | $r_{rb}$ | $p_{\text{BM}}$ | AUC |
| --- | --- | --- | --- | --- | --- | --- | --- | --- | --- |
| CS-PRISM | HC vs UC | K2 Entropy | 34 | 50 | 1209 | 0.0011 | -0.422 | $3.74 \times 10^{-4}$ | 0.711 |
| CS-PRISM | HC vs UC | MPA Entropy | 34 | 50 | 1225 | $6.43 \times 10^{-4}$ | -0.441 | $1.38 \times 10^{-4}$ | 0.721 |
| CS-PRISM | HC vs UC | GSL $\log(E)$ | 34 | 50 | 1278 | $9.79 \times 10^{-5}$ | -0.504 | $1.48 \times 10^{-5}$ | 0.752 |
| CS-PRISM | HC vs CD | K2 Entropy | 34 | 67 | 1938 | $9.56 \times 10^{-9}$ | -0.701 | $4.48 \times 10^{-13}$ | 0.851 |
| CS-PRISM | HC vs CD | MPA Entropy | 34 | 67 | 1794 | $2.56 \times 10^{-6}$ | -0.575 | $4.33 \times 10^{-9}$ | 0.788 |
| CS-PRISM | HC vs CD | GSL $\log(E)$ | 34 | 67 | 2040 | $9.71 \times 10^{-11}$ | -0.791 | $2.37 \times 10^{-22}$ | 0.896 |
| LSS-PRISM | HC vs UC | K2 Entropy | 20 | 48 | 584 | 0.1636 | -0.217 | 0.2139 | 0.608 |
| LSS-PRISM | HC vs UC | MPA Entropy | 20 | 48 | 615 | 0.0702 | -0.281 | 0.0703 | 0.641 |
| LSS-PRISM | HC vs UC | GSL $\log(E)$ | 20 | 48 | 327 | 0.0401 | +0.319 | 0.0155 | 0.659 |
| LSS-PRISM | HC vs CD | K2 Entropy | 20 | 97 | 1030 | 0.6666 | -0.062 | 0.7190 | 0.531 |
| LSS-PRISM | HC vs CD | MPA Entropy | 20 | 97 | 1052 | 0.5551 | -0.085 | 0.6101 | 0.542 |
| LSS-PRISM | HC vs CD | GSL $\log(E)$ | 20 | 97 | 482 | $4.16 \times 10^{-4}$ | +0.503 | $7.34 \times 10^{-7}$ | 0.752 |
| Stinki | HC vs UC | K2 Entropy | 51 | 36 | 1446 | $5.47 \times 10^{-6}$ | -0.575 | $3.66 \times 10^{-7}$ | 0.788 |
| Stinki | HC vs UC | MPA Entropy | 51 | 36 | 1428 | $1.13 \times 10^{-5}$ | -0.556 | $6.53 \times 10^{-7}$ | 0.778 |
| Stinki | HC vs UC | GSL $\log(E)$ | 51 | 36 | 1602 | $3.85 \times 10^{-9}$ | -0.745 | $4.49 \times 10^{-15}$ | 0.873 |

